# Machine Learning-GWAS reveals the role of *WSD1* gene for cuticular wax ester biosynthesis and key genomic regions controlling early maturity in bread wheat

**DOI:** 10.1101/2023.11.03.565125

**Authors:** Honoré Tekeu, Martine Jean, Eddy L. M. Ngonkeu, François Belzile

## Abstract

This study employed Machine Learning-Genome-Wide Association Study (ML-GWAS) to identify genomic regions linked to cuticular wax ester biosynthesis (SW) and early maturity (DM) in wheat. Using a dataset with 170 wheat accessions and 74K SNPs, four GWAS tools (MLM, CMLM, FarmCPU, and BLINK) and five machine learning techniques (RF, ANN, SVR, CNN, and SVM) were applied. A highly significant SW association was found on chromosome 1A, with the peak SNP (chr1A:556842331) explaining 50% of the phenotypic variation. A promising candidate gene, *TraesCS1A01G385500*, was identified as an ortholog of Arabidopsis thaliana’s *WSD1* gene, which plays a crucial role in very long-chain (VLC) wax ester biosynthesis. For DM, four QTLs were detected on chromosomes 4B (two QTLs), 2A, and 5A. Haplotype analysis revealed that alleles TT significantly contribute to cuticular wax ester biosynthesis and early maturity in wheat varieties. The study underscores the superior performance of ML models, especially when combined with advanced multi-locus GWAS models like BLINK and FarmCPU, with significantly lower p-values for identifying relevant QTLs compared to traditional methods. ML approaches hold potential for revolutionizing the study of complex genetic traits, offering insights to enhance wheat crops’ resilience and quality. ML-GWAS emerges as a compelling tool for genomic-based breeding, enabling breeders to develop improved wheat varieties with greater precision and efficiency.

## Introduction

Wheat (*Triticum aestivum* L.), a globally essential staple crop, faces a multitude of environmental challenges, from drought and salinity to extreme temperatures and pest pressures (He et al., 2022). A vital component of its adaptive response to these challenges is the hydrophobic cuticle, composed of cutin and cuticular waxes (Wang and Chang, 2022). This lipidic shield defends against non-stomatal transpiration, UV radiation, pathogens, and insect invasions while maintaining the integrity of adjacent plant organs (Ingram and Nawrath, 2017; Martin and Rose, 2014).

The cuticle consists of two primary constituents: cutin, an insoluble polyester, and cuticular waxes, encompassing very-long-chain (VLC) fatty acids, aldehydes, ketones, esters, alcohols, alkanes, and other compounds (Kunst and Samuels, 2009). Alkanes, a significant component of cuticular waxes, play a critical role in enhancing plant drought tolerance (Kosma et al., 2009; Seo et al., 2011). In the realm of wheat, genes related to wax biosynthesis, including TaFARs for primary alcohols and the W1 locus for β-diketones, have been identified (Hen-Avivi et al., 2016; Y. Wang et al., 2015a, 2015b). One pivotal gene, TaCER1-1A, has been recognized for its involvement in alkane accumulation in wheat (Li et al., 2019). In a recent study by (He et al., 2022), attention is drawn to TaCER1-6A, another key gene involved in alkane biosynthesis in wheat, with investigations extending to overexpression and CRISPR/Cas9-mediated gene editing.

To date, no study has pinpointed a gene responsible for the biosynthesis of wax VLC esters, which play a crucial role in mitigating leaf water loss, particularly under drought conditions. The journey of these wax constituents from the Golgi and trans-Golgi network (TGN) to the plasma membrane and onward to the cuticle involves pathways coordinated by ABCG subfamily half transporters and lipid transfer proteins (LTPs) (DeBono et al., 2009; Ichino and Yazaki, 2022; Pighin et al., 2004; Wang and Chang, 2022).

Additionally, the cultivation of early-maturing wheat varieties holds critical importance in regions characterized by short growing seasons and extended daylight, exemplified by the Northern Great Plains of Canada and the USA (A. Kamran et al., 2013). Early maturation not only enhances crop yields but also acts as a safeguard against frost damage, a threat that can significantly compromise grain quality and overall agricultural productivity (Iqbal et al., 2007). The precise timing of wheat’s flowering is intricately regulated by a complex interplay of genes that dictate growth patterns and earliness. These genetic regulators encompass vernalization (Vrn), photoperiod (Ppd), and earliness per se (Eps) genes, shaping when wheat plants initiate flowering and influencing their growth habits (Atif Kamran et al., 2013).

Adding to this complexity, certain genetic factors, such as dwarfing genes, subtly affect the timing of heading, flowering, and maturity, introducing further intricacies in the regulation of these vital agricultural traits (Chen et al., 2018; Daoura Goudia et al., 2014). Earliness per se genes also play a role in enhancing the adaptability of wheat plants, contributing to their resilience in varying environments (Snape et al., 2001). Recent studies, including one by Semagn et al. (2021), have delved into the intricate genetic mapping of Quantitative Trait Loci (QTLs) associated with days to maturity, particularly in wheat varieties evaluated under both conventional and organic farming practices. These studies have identified key QTLs on chromosome 4B, shedding light on the genetic mechanisms governing maturity. Furthermore, earlier research by authors such as (Zou et al., 2017a, 2017b) employing extensive genetic mapping using 1203 markers in RIL populations like ‘Attila’ and ‘CDC Go’ has uncovered a shared genomic region linked to maturity, situated on both chromosome 4B and 5A.

Among the array of genetic factors at play, certain dwarfing genes, including Rht-B1, Rht5, Rht8, and Rht12, have been identified as contributing factors, subtly influencing the timing of heading, flowering, and maturity in wheat varieties. These genetic elements add an additional layer of complexity to the intricate regulation of these pivotal traits (Chen et al., 2018; Daoura Goudia et al., 2014).

While molecular markers have facilitated characterizing genetic diversity, phenotypic assessments have primarily determined the utility of these genetic resources in breeding (Belzile et al., 2020). With the availability of high-density SNP markers, Genome-Wide Association Studies (GWAS) have become a powerful tool for identifying and mapping loci contributing to phenotypic variation among diverse genetic materials that have undergone extensive recombination (Yu and Buckler, 2006). Recent applications of highly reproducible GBS-derived SNPs have uncovered candidate genes influencing grain size in bread wheat (Tekeu et al., 2021). GWAS has become a standard approach across species for identifying genes associated with critical traits (Ashkenazy et al., 2022).

However, there remain challenges with conventional GWAS techniques, including the “large p, small n” issue when the number of markers surpasses the number of genotypes (Kaler et al., 2020; Mohammadi et al., 2020). Conventional GWAS methods are better suited for identifying common SNPs with substantial main effects, while the distinction between causal variants and correlated genes linked by linkage disequilibrium remains problematic (Enoma et al., 2022; Nicholls et al., 2020). Moreover, conventional GWAS approaches lack the power to uncover minor-effect SNPs associated with specific traits (Zhou et al., 2019). Consequently, machine learning (ML) techniques offer an opportunity to address these limitations and gain insights into the complex genetic basis of traits, as demonstrated in other crop species (Ashkenazy et al., 2022; Kwon et al., 2022).

Machine learning models for GWAS vary in complexity, from simple logistic regression to sophisticated ensemble models such as random forests, gradient boosting, and neural networks. These ML algorithms focus on maximizing prediction accuracy and excel at capturing multi-locus SNP interactions better than conventional methods. Support Vector Regression (SVR) is one such machine learning technique that has shown promise in predicting important agricultural traits (Yoosefzadeh Najafabadi et al., 2021). While SVR has found application in various crop studies, the potential of other ML techniques, such as Random Forest (RF), Convolutional Neural Networks (CNN), Artificial Neural Networks (ANN), and Support Vector Machines (SVM), remains largely untapped when compared to the more conventional GWAS tools like Mixed Linear Model (MLM), Compressed Mixed Linear Model (CMLM), Fixed and random model Circulating Probability Unification (FarmCPU), and Bayesian-information and Linkage-disequilibrium Iteratively Nested Keyway (BLINK).

This study aims to address this gap and provide valuable insights to bolster crop resilience. By employing a diverse array of advanced ML techniques in GWAS analysis, we seek to identify the genomic regions associated with cuticular wax ester biosynthesis (SW) and early maturity (DM) in wheat. Our approach promises to shed light on the intricate genetic mechanisms governing these vital traits and contribute to the advancement of crop breeding efforts for improved wheat varieties. Our research hypotheses revolve around specific genomic regions influencing SW and DM in a diverse global collection of bread wheat accessions. Furthermore, it postulates that ML-GWAS approaches will outperform traditional GWAS methods, in identifying Quantitative Trait Loci (QTLs) relevant to SW and DM traits in wheat. The present study aims to decipher the genetic underpinnings of SW and DM using ML-GWAS approaches.

## Materials and methods

### Plant materials

In this study, an international collection of 170 accessions was employed for genome-wide association analyses. These cultivars were obtained from various international wheat breeding programs. The South African accessions consisted of spring wheat lines from the Western Cape region, along with some winter bread wheat lines from other parts of the country. The East African spring-type accessions were gathered in Kenya and Ethiopia. The Mexican accessions were obtained via the International Maize and Wheat Improvement Center (CIMMYT), and they included spring accessions from Mexicali and Baja California. The Central African accessions were provided by the Institute of Agricultural Research for Development (IRAD) and farmers (Tekeu et al., 2017). The French accessions were winter lines, and those from North Africa were composed of spring lines acquired from the International Center for Agricultural Research in the Dry Areas (ICARDA).

### Phenotyping

A panel of 170 accessions of bread wheat was phenotyped and used for genome-wide association analyses. Field trials were conducted in two different locations in the bimodal humid forest zone of Cameroon, during the 2015-2016 season in Munt Mbankolo (1057 m above sea level) and during 2016-2017 in Nkolbisson (650 m a. s. l.). At each trial site, an incomplete alpha-lattice design with two replications was used and each accession was planted, as previously reported by (Tekeu et al., 2021). Then, fields trials were managed in accordance with the technical recommendations and standard agricultural practices for wheat (Pask et al., 2012). Spike waxiness (SW; 0: Absent, 2: Almost none, 3: Very little, 4: Little, 5: Intermediate, 6: Some, 7: Much, 8: Very much) and DM (days-to-maturity) were assessed when 50% of spikes had turned yellow (Zadoks et al., 1974).

### Analysis of phenotypic data

We conducted the analysis of variance for each trait using PROC MIXED in SAS 9.4. In this analysis, each cultivar was considered a fixed effect, while replications and environments were treated as random effects. Pearson correlation coefficients between pairs of phenotypic traits were computed using Pearson’s correlation in SPSS 20.0. To assess the heritability of each trait, we utilized the broad-sense heritability (h²) formula: h² = VG / (VG + VGE + Ve), where VG represents genetic variance, VGE is the genetic-environment interaction variance, and Ve is the error variance.

### DNA isolation, GBS library construction and sequencing

To extract genomic DNA from dried young leaf tissue (∼ 5 mg) of all accessions, we used a CTAB DNA isolation method (Doyle and Doyle, 1990). The extracted DNA was quantified using a Quant-iT™ PicoGreen kit (ThermoFisher Scientific, Canada), and concentrations were normalized to 20 ng/μl for library preparation. We constructed three 96-plex *PstI-MspI* GBS libraries as described by (Elshire et al., 2011). Subsequently, each library was sequenced on three P1 chips using an Ion Torrent PGM sequencer at the Plate-forme d’Analyses Génomiques of the Institut de Biologie Intégrative et des Systèmes (Université Laval, Québec, Canada).

### Single nucleotide polymorphism calling and bioinformatics analysis

Genomic DNA sequences of wheat samples, with an average of 2.4 million reads per wheat line, were analyzed using the FastGBS pipeline (Torkamaneh et al., 2017). The reads were aligned to the wheat reference genome (Chinese Spring v1.0), and SNPs were called using FastGBS. Standard filtration steps were applied to the FastGBS results, as previously described by (Tekeu et al., 2021). Additional filtration steps were carried out on this subset to retain only SNPs with a minor allele frequency (MAF) of at least 0.05.

### Machine Learning-Genome-Wide Association Study

We conducted a genome-wide association study (GWAS) to identify genomic regions associated with variation in SW and DM using a dataset comprising 170 accessions and 74K single nucleotide polymorphisms (SNPs). We employed a comprehensive approach that integrated four GWAS analytical methods, namely the Mixed Linear Model (MLM), Compressed Mixed Linear Model (CMLM), Fixed and random model Circulating Probability Unification (FarmCPU), and Bayesian-information and Linkage-disequilibrium Iteratively Nested Keyway (BLINK). In addition, we harnessed the power of five machine learning algorithms, which included Random Forest (RF), Support Vector Regression (SVR), Convolutional Neural Networks (CNN), Artificial Neural Networks (ANN), and Support Vector Machines (SVM). This integrated approach allowed us to assess the association between SNP markers and estimated genotypic values (BLUEs) for each trait.

For MLM, CMLM, FarmCPU, and BLINK methods, we made use of the Genomic Association and Prediction Integrated Tool (GAPIT) version 2 (Lipka et al., 2012) in conjunction with the rMVP packages (Yin et al., 2021). Our association analyses were performed while correcting for both population structure and relationships among individuals, with the incorporation of either the Q+K matrices. The K matrix was computed using the Van Raden method (Lipka et al., 2012). The significance threshold for genome-wide association was determined based on a false discovery rate (FDR-adjusted p < 0.05).

In the case of machine learning algorithms, we utilized a scaled method (ranging from 0 to 100) to estimate the importance of each SNP associated with the traits of interest. To integrate the machine learning approach into GWAS, we implemented a five-fold cross-validation strategy with ten repetitions to estimate the variable importance of each SNP, following (Siegmann and Jarmer, 2015). Therefore, we applied a global empirical threshold, as proposed by (Churchill and Doerge, 1994; Doerge and Churchill, 1996). This threshold was determined by fitting the ML algorithm, recording SNPs with the highest variable importance scores, repeating the process 1000 times, and selecting associated SNPs based on α = 0.5. The machine learning methods were executed using the Caret package (Kuhn et al., 2020) in R software version 4.2.2. Throughout these analyses, we ensured that association analysis was conducted while correcting for both population structure and relationships among individuals, using a combination of the Q + K matrices. The p-value threshold for significance in the genome-wide association was determined based on a false discovery rate (FDR-adjusted p ≤ 0.05).

### Identification of candidate genes and haplotype analysis

To identify candidate genes contributing to SW and DM, we defined haplotype blocks containing the peak SNP. Each region with the peak SNP was visually explored for its LD structure and for genes located in such regions, and the annotated genes within each interval were screened thanks to the annotated and ordered reference genome sequence in place by (International Wheat Genome Sequencing Consortium (IWGSC), 2018). Candidate genes potentially involved in each trait were further investigated. The function of these genes was also inferred by a BLAST of their sequences to the UniProt reference protein database (http://www.uniprot.org/blast/). To further provide more information about potential candidate genes, we used RNA-seq data of (Ramírez-González et al., 2018), based on the electronic fluorescent pictograph (eFP) at bar.utoronto.ca/eplant (by (Waese et al., 2017) to identify in what tissues and at which developmental stages candidate genes were expressed in wheat.

To better define the possible alleles in a strong candidate gene and trait, we defined haplotypes around the peak SNP. For each haplotype, we calculated the trait mean for lines sharing the same haplotype using the R ggpubr program.

## Results

### Phenotypic characterization

In order to delve into the traits of SW and DM in wheat, we meticulously assessed their phenotypes over the span of two years at two distinct sites. As summarized in Table 1, the observed means (± standard deviation) for these traits were as follows: 5.35 (±1.56) for SW and 98.06 days (±4.65) for DM. The broad-sense heritability estimates were robust, measuring 55.4% for SW and 50% for DM. An analysis of variance uncovered noteworthy differences attributable to genotypes (G) for all traits, and, in the case of SW and DM, the interaction between genotype and environment (GxE) also emerged as a significant factor. A correlation analysis unveiled a highly significant positive correlation between SW and DM (r = 0.273; p < 0.01).

**Table 1.**
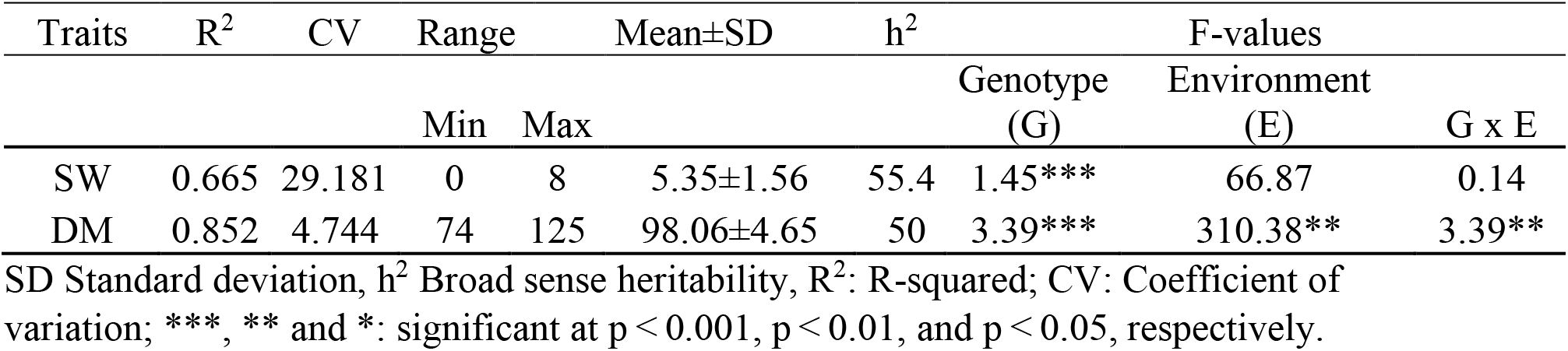
Descriptive statistics, broad-sense heritability (h²), and F-values from variance analysis for two key agronomic traits in a cohort of 170 wheat lines.

| Traits | $R^2$ | CV | Range | | Mean±SD | $h^2$ | F-values | | |
| --- | --- | --- | --- | --- | --- | --- | --- | --- | --- |
|  |  |  | Min | Max |  |  | Genotype (G) | Environment (E) | G x E |
| SW | 0.665 | 29.181 | 0 | 8 | 5.35±1.56 | 55.4 | 1.45*** | 66.87 | 0.14 |
| DM | 0.852 | 4.744 | 74 | 125 | 98.06±4.65 | 50 | 3.39*** | 310.38** | 3.39** |

SD Standard deviation, $h^2$ Broad sense heritability, $R^2$ : R-squared; CV: Coefficient of variation; \*\*\*, \*\* and \*: significant at $p < 0.001$ , $p < 0.01$ , and $p < 0.05$ , respectively.

Upon scrutinizing the relationship between SW and DM using bagplots analysis with the 170 accessions in our collection, no outliers were detected when considering the interplay between these two traits (Supplementary Figure S1). Consequently, for subsequent analyses including those involving population structure and genome-wide association studies (GWAS), all accessions were retained. The distribution of phenotypic traits appeared to approximate a normal distribution and exhibited characteristics of quantitative inheritance (Figure 1).

**Figure 1.**
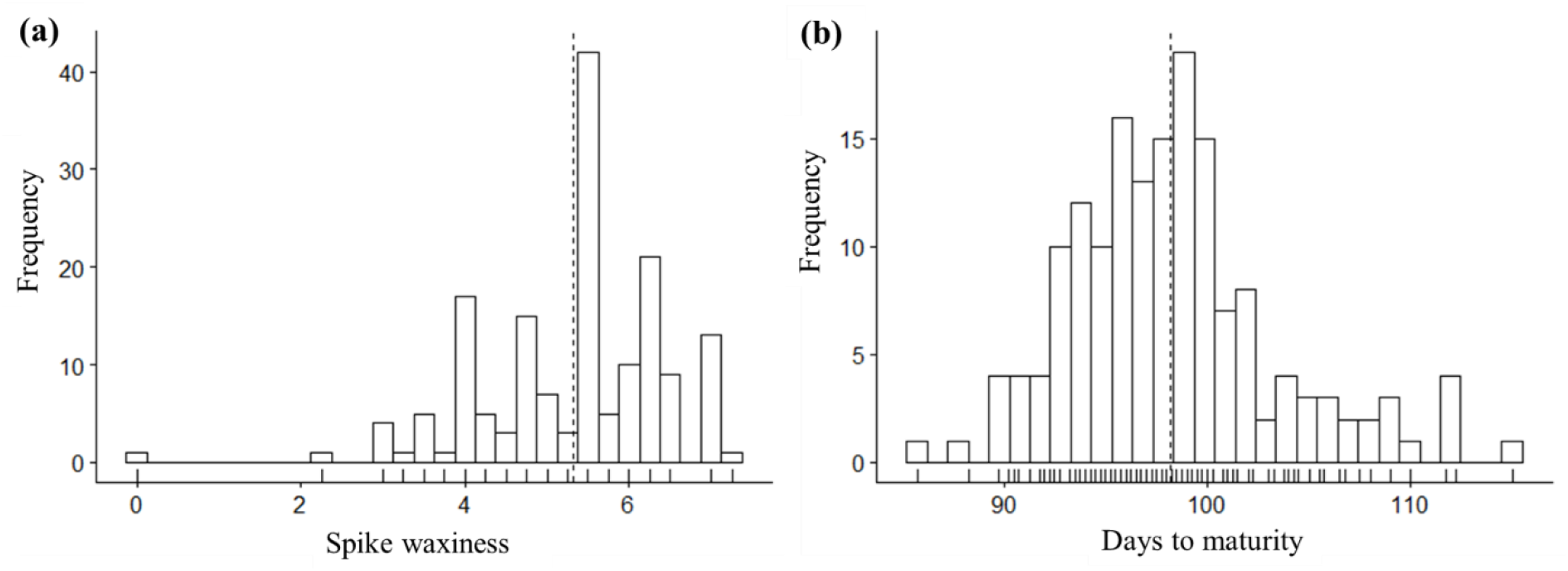
Distribution of phenotypes for spike waxiness (a) and days-to-maturity (b). Histograms are based on the average trait value of each wheat line across the different environments. The bars under the histograms represent the density of individuals.

### Genome coverage and population structure

Our comprehensive analysis revealed a total of 73,784 polymorphic SNP markers that spanned across the 21 chromosomes of the wheat genome, as depicted in Figure 2. As previously reported in our prior study, the examination of population structure within the accessions of this association panel revealed that K=6 provided the optimal representation of population structure within this set of accessions. These clusters notably aligned with the geographic regions of origin. The distribution of wheat accessions among these six subpopulations ranged from 6 to 43, with the largest number of accessions hailing from northwestern Baja California, Mexico, specifically represented by Mexico 1 (43). Conversely, the smallest subpopulation was observed in East and Central Africa, encompassing just 6 accessions.

**Fig 2.**
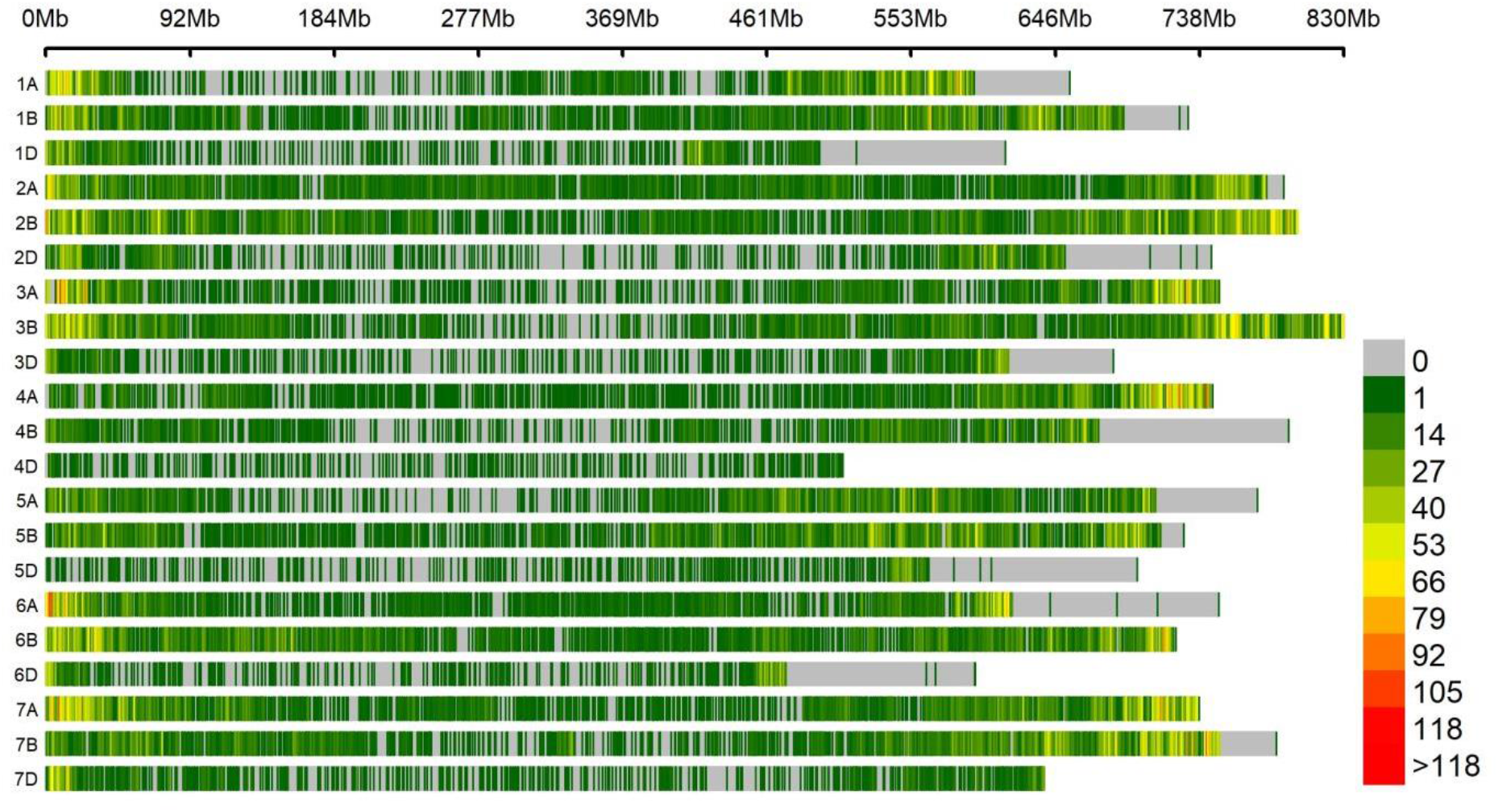
Genome coverage of polymorphic SNP markers over the physical map of the 21 chromosomes of the hexaploid wheat lines. The color reflects the density of SNP markers (i.e. number of SNPs within a sliding 1-Mb window).

### Marker-trait associations

To uncover the genomic regions responsible for the variation in SW and DM, we conducted an association analysis (GWAS) on a subset of accessions with phenotypic data (170 accessions and 73,784 SNPs). In this analysis, we employed four GWAS analytical tools (MLM, CMLM, FarmCPU, and BLINK), complemented by five machine learning techniques (RF, ANN, SVR, CNN, and SVM). Notably, the quantile-quantile (QQ) plots in Figure 3 demonstrated the effective control of confounding effects related to population structure and relatedness by all conventional GWAS and machine learning models. Deviations from the diagonal were observed only for the most extreme p-values, indicating a well-controlled analysis for both traits.

**Figure 3.**
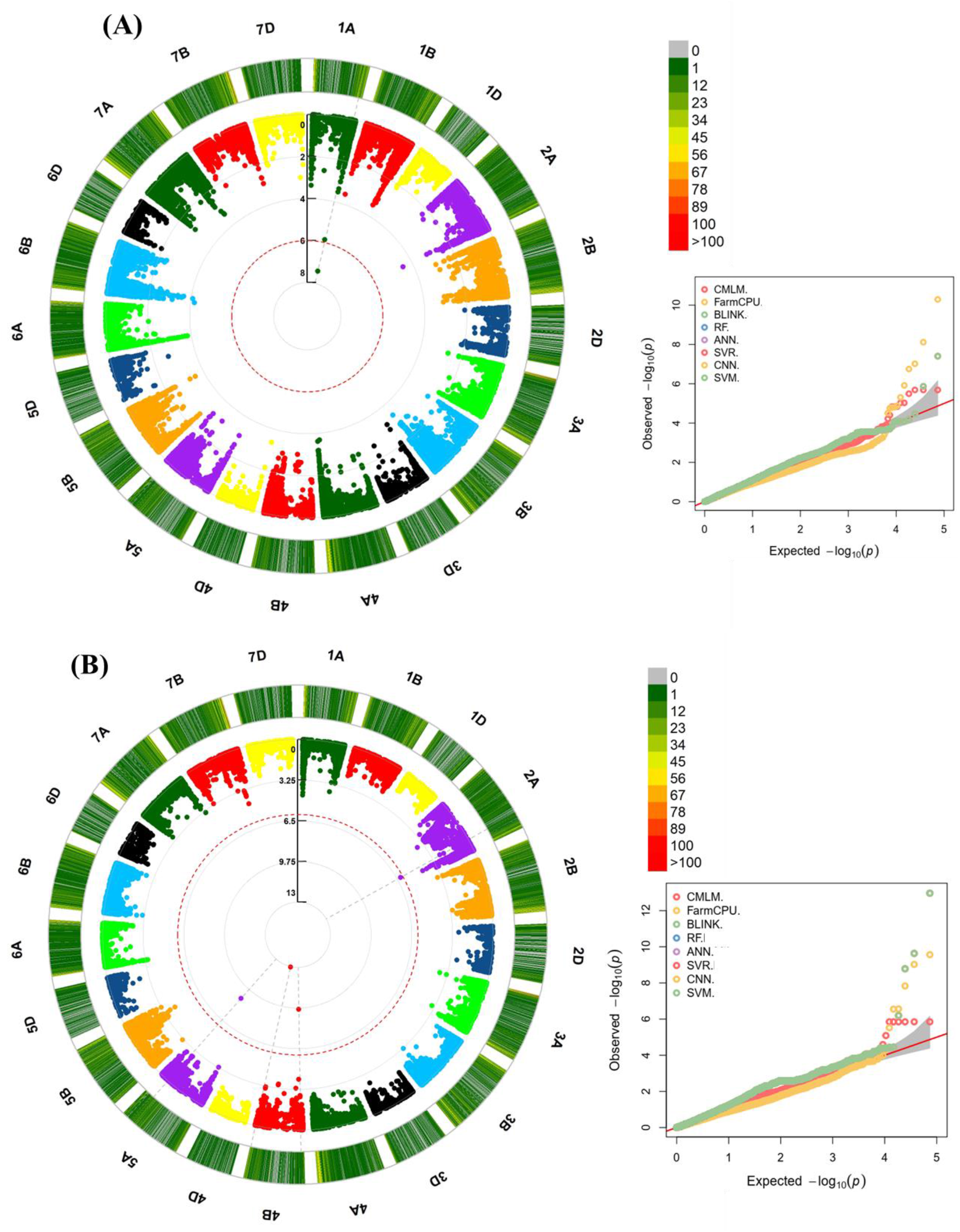
Genome-wide association analysis of 170 hexaploid wheat cultivars. Manhattan and Q-Q plots for all models shows the degree of association between SNPs and SW (A) and DM (B). In both cases, associations are declared significant at an FDR ⩽0.05. One marker (see the red circle) displayed significant associations with the SW trait. Four SNP markers (see the red circle) displayed significant associations with the DM.

For the SW trait, the results of the association analyses are visualized in the Manhattan plots presented in Figure 3. Using a threshold for false discovery rate (FDR) of ≤ 0.05 (as detailed in Supplementary Figure S2, marked by the green horizontal line), we identified four QTLs. Remarkably, only one QTL was co-identified by at least two models (Figure 4). The most robust and consistent association, located on chromosome 1A, is summarized in Table 2.

**Figure 4.**
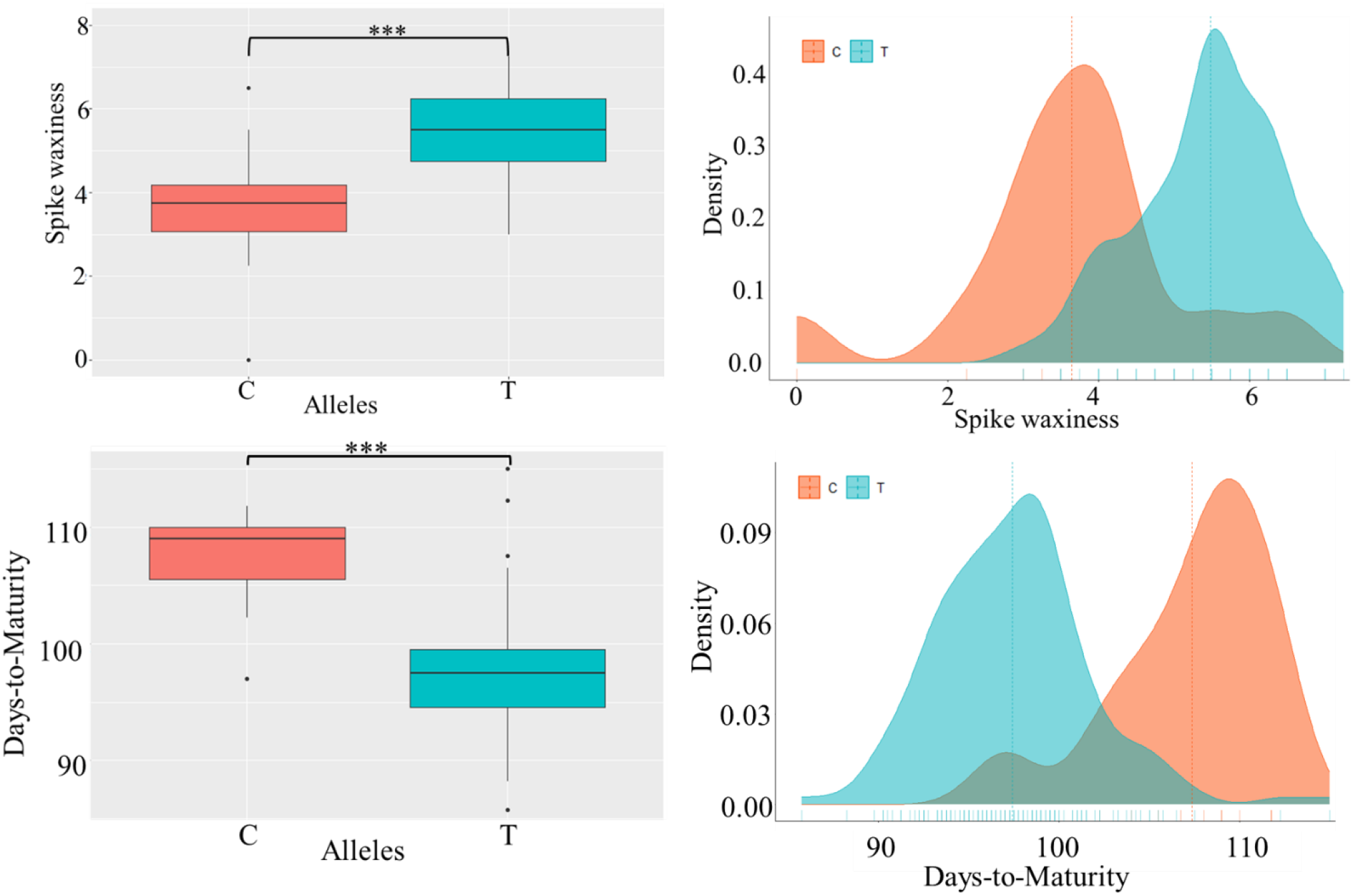
Boxplots (lef) and bimodal distribution (right) of Spike Waxiness and Days-to-maturity are represented for each haplotype. ***: signifcant at *P* < 0.001

**Table 2.**
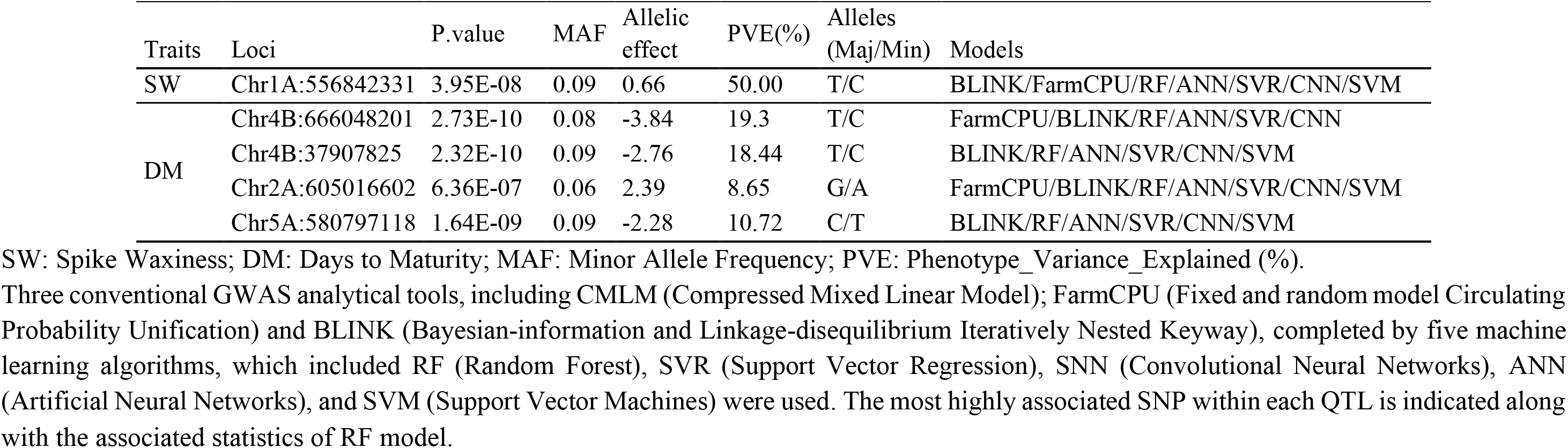
Details of loci associated with phenotypic traits identified by at least two methods in wheat.

This particular QTL was defined by its peak SNP, marked as chr1A:556842331, and was identified by the multi-locus models (FarmCPU and BLINK) as well as all five machine learning algorithms (RF, ANN, SVR, CNN, and SVM). Notably, this QTL explained a substantial 50% of the phenotypic variation observed in SW. The minor allele frequency (MAF) at this locus was 0.09, and it exhibited an allelic effect of 0.66.

These findings highlight a significant and consistent genetic association with SW on chromosome 1A, showcasing the power of both traditional GWAS and machine learning approaches in identifying key genomic regions influencing this trait.

Turning our attention to the DM trait, our investigation unveiled a total of eight genomic regions that displayed significant associations. The results of these association analyses are visualized in the Manhattan plots featured in Figure 3, with a stringent threshold for false discovery rate (FDR) of ≤ 0.05, as outlined in Supplementary Figure S2 and highlighted by the green horizontal line. However, we noted the co-identification of only four Quantitative Trait Loci (QTLs) by at least two models (Figure 3). Among these, the most robust associations, localized on chromosomes 4B, 5A, and 2A, are thoughtfully summarized in Table 2.

Of noteworthy mention is chr4B:666048201, which emerged as the peak SNP and was jointly identified by both multi-locus GWAS models (FarmCPU and BLINK) and four machine learning algorithms (RF, ANN, SVR, and CNN). These markers formed a robust linkage block, with all markers exhibiting perfect linkage disequilibrium (LD) (r^2^ = 1), as detailed in Supplementary Table S1. This discovery delineated a single QTL, with the peak SNP accounting for a substantial 19.3% of the phenotypic variation associated with DM. The minor allele frequency (MAF) at this locus was observed to be 0.08, while the allelic effect amounted to 3.84 days (Table 2).

In addition, another noteworthy association with DM on chromosome 4B was unveiled, defined by the peak SNP chr4B:37907825. This association was identified by the GWAS model BLINK and all five machine learning methods (RF, ANN, SVR, CNN, and SVM). It explained 18.44% of the phenotypic variation for DM, with a MAF of 0.09 and an allelic effect of −2.76 days.

Moreover, a QTL residing on chromosome 2A was brought to light, marked by the peak SNP chr2A:605016602, which explained 8.65% of the phenotypic variation for DM. This QTL was detected using both multi-locus GWAS models (FarmCPU and BLINK) and all five machine learning methods (RF, ANN, SVR, CNN, and SVM).

Furthermore, an additional QTL on chromosome 5A, characterized by the peak SNP chr5A:580797118, was identified through the BLINK model and all five machine learning methods (RF, ANN, SVR, CNN, and SVM). This QTL contributed to 0.72% of the phenotypic variation associated with DM.

These findings underscore the efficacy of our approach in uncovering key genomic regions associated with DM and highlight the potential of both traditional GWAS and machine learning techniques in unraveling the genetic underpinnings of complex traits.

Overall, the GWAS and ML methods successfully mitigated the confounding effects of population structure and relatedness and identified multiple genomic regions associated with spike waxiness and Days to maturity in wheat. These findings can provide insights into the genetic architecture of these traits and aid plant breeders in developing new bread wheat varieties with improved SW and maturity.

In order to gain a deeper understanding of the relationship between the peak SNP (chr1A:556842331) and SW, we delved into the realm of SNP haplotypes. Through a thorough analysis of haplotypes encompassing this peak marker, we unveiled two distinct haplotypes (Figure 4). Remarkably, we observed a notable divergence in phenotypic outcomes between these haplotypes. Haplotype TT displayed significantly higher values (5.481) compared to the values associated with haplotype CC (3.642). This revelation suggests that SNP markers flanking this gene could serve as valuable tools in marker-assisted breeding programs. By selecting alleles that contribute to drought-resistant wheat varieties, these programs hold the potential to enhance wheat productivity and bolster its resilience in the face of water scarcity.

To further refine our understanding of the association between the peak SNP (chr4B:666048201) and DM, we embarked on an exploration of SNP haplotypes. This investigation uncovered two distinct haplotypes encircling the peak SNP. Notably, our scrutiny of these haplotypes revealed a substantial difference in phenotypic outcomes (Figure 4). Haplotype TT was linked to significantly lower values (97.41) in comparison to haplotype CC (107.36). This observation posits that SNP markers flanking this gene have the potential to be valuable assets in marker-assisted breeding programs. By selecting alleles conducive to the development of short-season wheat varieties, these programs can contribute to the improvement of wheat productivity and the creation of cultivars better equipped to thrive in varying environmental conditions. More details are provided in Supplementary Table S2.

### Identification of candidate genes

To pinpoint the candidate genes that potentially govern SW and DM in our diverse wheat collection, we conducted an analysis of the genes located within the same linkage block as the peak SNP for each QTL.

In the genomic interval encompassing the QTL that contributes the most to the phenotypic variation in SW (50%) specifically, the region from 1A_555 to 557 Mb, surrounding the peak SNP (chr1A:556842331), we identified a total of 24 high-confidence genes. Upon a detailed examination of the gene annotations and expression profiles, one gene, TraesCS1A01G385500 on chromosome 1A, emerged as the most promising candidate. TraesCS1A01G385500 is an ortholog of the *Arabidopsis Thaliana* O-acyltransferase gene, commonly known as WSD1, a bifunctional wax ester synthase/diacylglycerol acyltransferase, involved in cuticular wax biosynthesis and essential for reducing leaf water loss, particularly during drought conditions. WSD1 has also been associated with the biosynthesis of very long-chain (VLC) wax esters, contributing to drought tolerance in Arabidopsis. This gene exhibits the highest expression levels in spike, roots, and shoot axis tissues (Figure 5). More details are provided in Supplementary Table S3.

**Figure 5.**
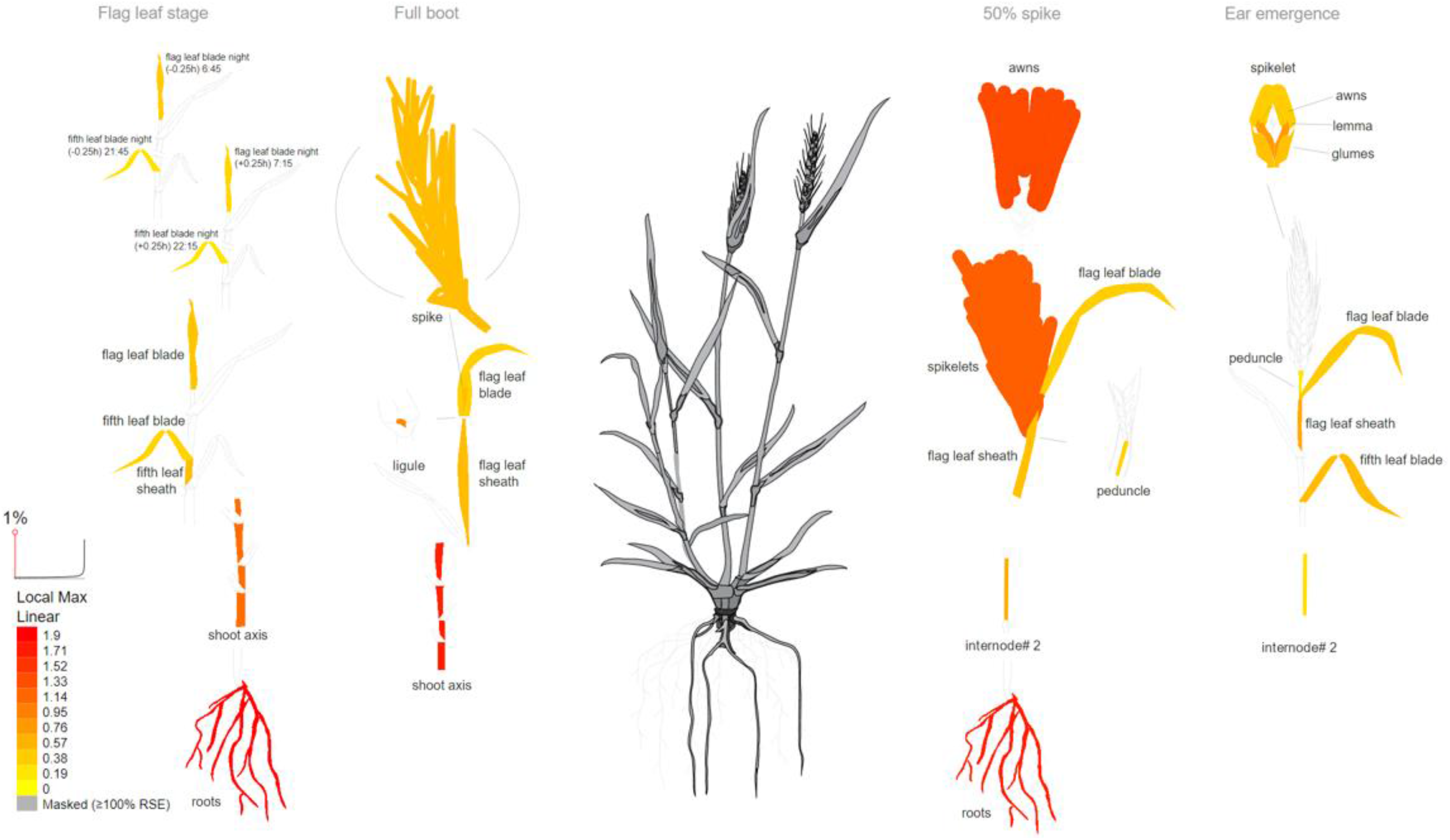
Expression profile of *TraesCS1A01G385500* gene based on transcriptomic analysis in wheat. As shown, this gene is most expressed in spike, roots and shoot axis and the image was generated with the eFP (RNA-Seq data) at http://bar.utoronto.ca/eplant/ by Waese et al.51. The legend at bottom lef presents the expression levels, coded by colors (yellow=low, red=high).

In our quest to identify potential candidate genes governing DM trait in our diverse wheat collection, we performed a meticulous analysis of the genes residing within the same linkage block as the peak SNP for each QTL. Within the genomic interval encompassing the QTL that makes the most substantial contribution to the phenotypic variation in DM (comprising 19.3% of the total variation), specifically spanning from 4B_666 to 668 Mb and surrounding the peak SNP (chr4B:666048201), we pinpointed a total of 27 high-confidence genes. Through an in-depth examination of gene annotations and expression profiles, no one emerged as the most promising candidate. More details are provided in Supplementary Table S3.

### Comparison of ML and conventional GWAS methods for identifying genomic regions

In our study, the peak SNP position on chr1A:556842331 exhibited the highest allelic effect (0.66) and explained a substantial phenotypic variance (50%) among all the identified SNPs associated with wheat SW trait. These associations were successfully detected using both GWAS models (BLINK and FarmCPU) and five machine learning algorithms (RF, ANN, SVR, CNN, and SVM). Importantly, conventional methods (MLM and CMLM) failed to identify this peak SNP linked to the SW trait (Supplementary Table S4). Furthermore, the machine learning models produced significantly lower p-values (CNN, RF, SVM, SVR, and ANN, all with p-values of 3.95E-08) compared to conventional GWAS models (CMLM with a p-value of 5.97E-05, MLM with a p-value of 5.97E-05, FarmCPU with a p-value of 6.34E-03, and BLINK with a p-value of 3.95E-08).

Shifting our focus to the DM trait, our investigation revealed the co-identification of four QTLs on chromosomes 4B, 5A, and 2A (with peak SNPs chr4B:666048201, chr4B:37907825, chr5A:580797118, and chr2A:605016602) by both GWAS models (BLINK and FarmCPU) and the five machine learning algorithms (RF, ANN, SVR, CNN, and SVM). The details of these robust associations are succinctly presented in Table 2. Regrettably, conventional methods (MLM and CMLM) were unable to detect these SNP peaks associated with the DM trait. Additionally, the machine learning models yielded significantly lower p-values for each of the associated markers compared to conventional GWAS models (Supplementary Table S4).

For the SNP chr4B:666048201, the machine learning models generated markedly lower p-values (CNN, RF, SVM, SVR, and ANN, all with p-values of 1.09E-13) compared to conventional GWAS models (CMLM with a p-value of 1.42E-06, MLM with a p-value of 1.42E-06, FarmCPU with a p-value of 2.73E-10, and BLINK with a p-value of 1.09E-13).

For the SNP chr4B:37907825, the machine learning models produced substantially lower p-values (CNN, RF, SVM, SVR, and ANN, all with p-values of 2.32E-10) compared to conventional GWAS models (CMLM with a p-value of 2.65E-04, MLM with a p-value of 2.65E-04, FarmCPU with a p-value of 4.81E-04, and BLINK with a p-value of 2.32E-10).

For the SNP chr5A:580797118, the machine learning models yielded notably lower p-values (CNN, RF, SVM, SVR, and ANN, all with p-values of 1.64E-09) compared to conventional GWAS models (CMLM with a p-value of 4.72E-04, MLM with a p-value of 4.72E-04, and FarmCPU with a p-value of 1, and BLINK with a p-value of 1.64E-09).

For the SNP chr2A:605016602, the machine learning models generated significantly lower p-values (CNN, RF, SVM, SVR, and ANN, all with p-values of 1.64E-09) compared to conventional GWAS models (CMLM with a p-value of 1.38E-01, MLM with a p-value of 1.38E-01, and FarmCPU with a p-value of 8.63E-02, and BLINK with a p-value of 6.36E-07).

In essence, these traditional methods used for genetic data analysis and the establishment of associations between genetic variations and traits have limitations when it comes to identifying subtle or minor effects of certain SNPs on SW and DM characteristics in wheat. On the contrary, machine learning algorithms, particularly in conjunction with the recent multi-locus GWAS models (BLINK and FarmCPU), exhibited superior performance in identifying relevant QTLs when compared to traditional MLM and CMLM methods.

## Discussion

In this study, we employed four GWAS models and five machine learning algorithms to investigate the genomic regions associated with spike waxiness and days to maturity within a dataset consisting of 170 accessions and 74K SNPs. Our analyses consistently identified a robust QTL located on chromosome 1A, demonstrating significance across both conventional GWAS models (FarmCPU and BLINK) and a variety of machine learning models (BLINK, RF, ANN, SVR, CNN, and SVM). Notably, the peak SNP (chr1A:556842331) within this QTL explained a substantial portion, 50%, of the phenotypic variation observed. Within the genomic interval encompassed by this QTL (1A_555 to 557 Mb) and centered around the peak SNP (chr1A:556842331), we identified a total of 24 high-confidence genes. Upon closer examination of gene annotations and expression profiles, one candidate gene, *TraesCS1A01G385500* on chromosome 1A, stood out as particularly promising. This gene exhibits high expression levels in spike, roots, and shoot axis tissues and shares orthology with the *Arabidopsis Thaliana* O-acyltransferase gene, widely known as the *WSD1* gene. Previous research has highlighted the significance of the *WSD1* gene, which serves as a bifunctional wax ester synthase/diacylglycerol acyltransferase (Li et al., 2008; Patwari et al., 2019). Its involvement in cuticular wax biosynthesis is well-documented, and it plays a pivotal role in reducing leaf water loss, particularly during drought conditions (Li et al., 2008; Patwari et al., 2019). The WSD1 gene has also been associated with the biosynthesis of very long chain (VLC) wax esters, contributing to drought tolerance in Arabidopsis (Patwari et al., 2019). VLC primary alcohols and acyl-CoAs serve as precursors for wax ester biosynthesis, catalyzed by the bifunctional wax ester synthase/diacylglycerol acyltransferase WSD1 (Li et al., 2008; Patwari et al., 2019). These wax components, including VLC fatty acids, aldehydes, alkanes, alcohols, ketones, and esters, undergo trafficking through the Golgi and trans-Golgi network (TGN) pathways to the plasma membrane (PM). From there, they are exported to the cuticle via ABCG subfamily half transporters and lipid transfer proteins (LTPs) (DeBono et al., 2009; Ichino and Yazaki, 2022; Pighin et al., 2004; Wang and Chang, 2022).

Moreover, prior studies have revealed the role of AtCER1 in VLC alkane biosynthesis in Arabidopsis (Aarts et al., 1995; Bourdenx et al., 2011; Sakuradani et al., 2013). Recently, He et al. (2022) identified a homologous gene of AtCER1 in wheat, named TaCER1-6A, which shares 55% amino acid identity with AtCER1. Similar to previously reported AtCER1 orthologs, including rice OsCER1 (Ni et al., 2018), wheat TaCER1-1A (Li et al., 2019), Brachypodium BdCER1-8 (Wu et al., 2019), cucumber CsCER1 (W. Wang et al., 2015), and P. pratensis PpCER1 (Wang et al., 2021), TaCER1-6A also contains three specific His-rich motifs essential for VLC alkane biosynthesis (Bernard et al., 2012). Therefore, (He et al., 2022) speculated that TaCER1-6A likely plays a similar role in VLC alkane biosynthesis in wheat. Notably, we observed that alleles associated with higher wax content were more prevalent in lines originating from East African spring-type accessions (Kenya and Ethiopia) and North Africa. These accessions primarily consist of spring lines cultivated in arid regions and were acquired from the International Center for Agricultural Research in the Dry Areas (ICARDA).

Ultimately, our study has unveiled a promising candidate gene, TraesCS1A01G385500, linked to spike waxiness, with implications for cuticular wax biosynthesis and its role in drought tolerance, as established in previous research. This discovery sheds light on the genetic mechanisms underpinning spike waxiness in bread wheat, offering valuable insights for future breeding and crop improvement efforts.

Regarding DM, we identified four strong genomic regions significantly associated with the trait on chromosomes 4B, 2A and 5A. Our results were consistent with those of (Semagn et al., 2021), who performed QTL mapping in four RIL populations evaluated under conventional and organic management systems and reported two QTLs associated with days to maturity on chromosome 4B (explaining 20.8% of the phenotypic variances), where one (QMat.dms-4B.2) at chr4B:569184188-599613837 is located on the extremity of long chromosome 4B arm with the peak SNP chr4B:666048201 (explaining 19.3% of the phenotypic variation) that was jointly identified by both multi-locus GWAS models and four ML algorithms (RF, ANN, SVR, and CNN) in the present study. QTL mapping conducted by previous authors (Zou et al., 2017a, 2017b) in the ‘Attila’ and ‘CDC Go’ RIL populations using genetic maps of 1203 markers identified a coincident genomic region associated with maturity under both management systems on chromosome 4B and 5A. We also identified QTLs on chromosomes 4B and 5A, with peak SNPs chr4B:666048201, chr4B:37907825, and chr5A:580797118 explaining 19.3%, 18.44%, and 10.72%, of the phenotypic variation, respectively. The favorable alleles for those QTLs on 4B and 5A were most originated from the accessions of North Africa, including spring lines (Attila) acquired from the International Center for Agricultural Research in the Dry Areas (ICARDA). Interesting, previous works also identified two QTLs for maturity on chromosome 4B (QMat.dms-4B) and chromosome 5A (QMat.dms-5A.2), which individually explained 15.9% and 14.0% of the phenotypic variance, respectively, and together accounted for 29.9% of the phenotypic variance across seven environments (Zou et al., 2017a, 2017b). The favorable alleles for QMat.dms-4B and QMat.dms-5A.2 originated from ‘Attila’ and ‘CDC Go’, respectively. (Chen et al., 2020) also identified a QTL associated with maturity on chromosome 4B (4B_s4991673-4B_d1258252) using a linkage map of 4439 markers produced by DArTseq technology and phenotype data from ‘Peace’ and ‘Carberry’ RIL populations assessed for two years under organic management and conventional systems, consistent with our results.

Our investigation into candidate genes associated with maturity in wheat led us to a genomic interval spanning the QTL that contributes significantly to the phenotypic variation in Days-to-Maturity (19.3% of the variation). This region, located between 4B_666 and 668 Mb and centered around the peak SNP (chr4B:666048201), contained a total of 27 high-confidence genes. Our findings align with prior research that has identified genomic regions on chromosome 4B associated with maturity and housing candidate genes related to flower-promoting factors. Notably, certain dwarfing genes, such as Rht-B1, Rht5, Rht8, and Rht12, have been reported to exert slight delays in heading, flowering, and/or maturity time in wheat. These genetic factors add complexity to the regulation of these traits (Chen et al., 2018; Daoura Goudia et al., 2014). The discovery in the present study contributes to our understanding of the genetic factors underpinning wheat maturity and sets the stage for future research aimed at elucidating the molecular mechanisms involved.

The results our study highlight the remarkable superiority of machine learning (ML) models in identifying significant genetic associations compared to traditional Genome-Wide Association Study (GWAS) methods, as demonstrated through substantially lower p-values. For SW, the peak SNP was efficiently identified by both GWAS models (BLINK and FarmCPU) and the five ML algorithms, emphasizing their robustness. Notably, the ML models, including CNN, RF, SVM, SVR, and ANN, produced significantly lower p-values (3.95E-08) compared to the traditional GWAS models, which had p-values ranging from 5.97E-05 to 6.34E-03. Traditional methods (MLM and CMLM) failed to detect this critical SNP, showcasing their limitations in capturing minor genetic effects. Shifting the focus to DM, the robust associations identified by GWAS models and ML algorithms demonstrated that conventional methods (MLM and CMLM) were less effective, failing to detect these essential SNP peaks. Once again, ML models consistently delivered significantly lower p-values, underscoring their increased sensitivity and accuracy in identifying genetic markers linked to DM. The differences in p-values were substantial, with ML models consistently outperforming the traditional GWAS methods.

These findings reveal that traditional GWAS methods face limitations in detecting minor genetic effects on SW and DM traits in wheat. Conversely, ML models, especially when coupled with advanced multi-locus GWAS models like BLINK and FarmCPU, exhibited a superior performance characterized by significantly lower p-values. This work demonstrates the potential of ML approaches to revolutionize the study of complex genetic traits, offering valuable insights for crop improvement and stress resilience in bread wheat. Our hypotheses (1) regarding the presence of specific genomic regions associated with SW and DM in a diverse global collection of bread wheat accessions and (2) the superior performance of Machine Learning-Genome-Wide Association Study (ML-GWAS) approaches over traditional GWAS methods in identifying relevant genomic regions associated with SW and DM traits in bread wheat have been confirmed. Our study has provided evidence that conventional GWAS approaches, such as MLM, and CMLM, lack the ability to effectively detect SNPs with minor effects underlying specific traits. In other words, these traditional methods used to analyze genetic data and establish associations between genetic variations and traits are not sensitive enough to identify subtle or minor effects of certain SNPs on the characteristics of SW and DM in wheat. These findings align with previous research conducted by (Yoosefzadeh-Najafabadi et al., 2023; Zhou et al., 2019), which also highlighted the limited power of conventional GWAS approaches in detecting SNPs with minor effects on specific traits.

However, our study has revealed the effectiveness of an alternative approach, utilizing machine learning algorithms in GWAS. By employing this method, we were able to overcome the limitations of traditional GWAS and more accurately identify SNPs with smaller yet significant effects on SW and DM traits in wheat. Additionally, the most robust associations identified by modern GWAS methodologies (BLINK and FarmCPU) were reaffirmed by machine learning techniques. These results are consistent with the studies conducted by (Yoosefzadeh-Najafabadi et al., 2023) and (Zhou et al., 2019), which compared SVR and RF algorithms, respectively, to conventional GWAS methods in soybean. They reported that machine learning algorithms are more accurate and sensitive in detecting subtle or minor effects of certain SNPs on traits of interest. Additionally, our study demonstrated the effectiveness of the new GWAS model, BLINK and FarmCPU, in accurately and efficiently detecting SNPs with smaller but significant effects on SW and DM traits in wheat. In fact, both real and simulated data analyses have shown that BLINK significantly improves statistical power compared to FarmCPU, while also reducing computing time (Huang et al., 2019).

The FarmCPU, developed by (Liu et al., 2016), represents an iterative method that addresses the issue of false positive control and confounding between testing markers and cofactors simultaneously. As FarmCPU tests markers in a fixed-effect model, it is computationally more efficient than methods that test markers in a random-effect model, such as MLM, CMLM, ECMLM, SUPER, and MLMM (Liu et al., 2016). Studies have demonstrated the order of the statistical power of these methods: BLINK > FarmCPU > CMLM > MLM (Huang et al., 2019; Liu et al., 2016; Zhang et al., 2010).

The utilization of machine learning algorithms (RF, ANN, SVR, CNN, and SVM), along with the recent multi-locus GWAS model, BLINK, and FarmCPU has enabled a more sensitive and precise identification of genetic factors influencing specific traits. This opens up new opportunities for wheat improvement and selection. Indeed, ML algorithms are focused on maximizing prediction accuracy at the individual subject level and have been shown to capture multi-locus SNP interactions better than univariate association studies (Okser et al., 2014, 2013). Additionally, ML techniques provide an opportunity to better understand multi-locus genetic variants and their interactions in predicting complex traits (Ashkenazy et al., 2022; Kwon et al., 2022). This approach provides a more sophisticated and reliable means of discovering genetic markers associated with SW and DM traits, which can have significant implications for agriculture, varietal selection, and understanding the genetic mechanisms governing crop characteristics.

Overall, both GWAS and machine learning methods have successfully addressed the confounding effects of population structure and relatedness, allowing us to identify multiple genomic regions associated with SW and DM traits in wheat. These findings shed light on the genetic architecture of these traits and offer valuable insights to plant breeders in their efforts to develop new bread wheat varieties with improved SW and DM.

## Conclusion

In this study, our primary objective was to identify the genomic regions associated with SW and DM using state-of-the-art Machine Learning-Genome-Wide Association Study (ML-GWAS) techniques. Our findings provide a deep understanding of the genetic landscape governing these critical traits, delivering valuable insights that can significantly inform wheat breeding and crop improvement strategies. Leveraging ML-GWAS, we successfully identified a robust QTL significantly associated with SW on chromosome 1A, represented by the peak SNPs chr1A:556842331, explaining an impressive 50% of the phenotypic variation. Additionally, we detected four strong genomic regions significantly associated with DM on chromosomes 4B, 2A, and 5A, employing the same cutting-edge methods. Notably, our study unveiled a candidate gene linked to the QTLs for SW. *TraesCS1A01G385500*, an ortholog of the *Arabidopsis Thaliana* O-acyltransferase gene *WSD1*, plays a pivotal role in cuticular wax biosynthesis. It is essential for reducing leaf water loss, particularly during drought conditions, and contributes to drought tolerance through the biosynthesis of very long-chain (VLC) wax esters. Our study also shows that, ML models, especially when coupled with advanced multi-locus GWAS models like BLINK and FarmCPU, exhibited a superior performance characterized by significantly lower p-values in identifying relevant QTLs compared to traditional methods like MLM and CMLM. This work demonstrates the potential of ML approaches to revolutionize the study of complex genetic traits, offering valuable insights for crop improvement and stress resilience in bread wheat. ML-GWAS emerges as a compelling approach for genomic-based breeding strategies, providing breeders with more accurate and efficient tools to develop improved wheat varieties. Our research significantly advances the precision and effectiveness of GWAS, emphasizing the importance of incorporating advanced computational methods into crop breeding studies. The insights into the genetic architecture of SW and DM traits in wheat offer essential knowledge for designing targeted crop improvement strategies. Moreover, the versatility and effectiveness demonstrated by the ML-GWAS approach extend its applicability beyond wheat and can be harnessed to address other crop traits, thus enhancing progress in crop genetics research and breeding efforts on a broader scale.

Overall, the integration of machine learning techniques with GWAS stands as a potent tool for dissecting complex traits in crop genetics research. The findings of our study hold great promise for the field of wheat breeding and crop improvement strategies, making substantial contributions to enhancing agricultural productivity and ensuring food security in the face of evolving global challenges.

## Additional Information

Supplementary information for this paper is available at:

## Acknowledgements

Authors are grateful to the International Maize and Wheat Improvement Center (CIMMYT), the International Center for Agricultural Research in the Dry Areas (ICARDA) and the Plant Breeding Laboratory (Department of Genetics, Stellenbosch University) for their technical supports and wheat varieties collection. We would like to thank Dr Wuletaw Tadesse, Mr Tsimi Patrick, Mr Charly Mam for their technical support. We are grateful to Dr Amina Abed and Dr Jérôme Laroche for their technical support during bioinformatics analyses.

