## Supplementary Table S1 for "Machine Learning-GWAS reveals the role of *WSD1* gene for cuticular wax ester biosynthesis and key genomic regions controlling early maturity in bread wheat" (from another version of this preprint: version 1, 2023-11-05)

**Supplementary Table S1. LD information for associated SNP pair**

LD information for SNP pair [ chr4B:666052086 chr4B:666052143 ]

R-sq = 1.000	D' = 1.000

Haplotype	Frequency	Expectation under LE

---------	---------	--------------------

GT	0.085	0.007

CT	0.000	0.078

GC	0.000	0.078

CC	0.915	0.837

In phase alleles are GT/CC

LD information for SNP pair [ chr4B:666052086 chr4B:666052224 ]

R-sq = 1.000	D' = 1.000

Haplotype	Frequency	Expectation under LE

---------	---------	--------------------

GT	0.085	0.007

CT	0.000	0.078

GC	0.000	0.078

CC	0.915	0.837

In phase alleles are GT/CC

Analysis finished: Sat Oct 10 13:42:15 2020

LD information for SNP pair [ chr4B:666052086 chr4B:666048201 ]

R-sq = 1.000	D' = 1.000

Haplotype	Frequency	Expectation under LE

---------	---------	--------------------

GC	0.085	0.007

CC	0.000	0.078

GT	0.000	0.078

CT	0.915	0.837

In phase alleles are GC/CT

Analysis finished: Sat Oct 10 13:43:21 2020

LD information for SNP pair [ chr4B:666052086 chr4B:666052168 ]

R-sq = 1.000	D' = 1.000

Haplotype	Frequency	Expectation under LE

---------	---------	--------------------

GA	0.085	0.007

CA	0.000	0.078

GG	0.000	0.078

CG	0.915	0.837

In phase alleles are GA/CG

Analysis finished: Sat Oct 10 13:44:12 2020

LD information for SNP pair [ chr4B:666052086 chr4B:666052212 ]

R-sq = 1.000	D' = 1.000

Haplotype	Frequency	Expectation under LE

---------	---------	--------------------

GA	0.085	0.007

CA	0.000	0.078

GC	0.000	0.078

CC	0.915	0.837

In phase alleles are GA/CC

LD information for SNP pair [ chr4B:666052086 chr4B:425658194 ]

R-sq = 0.039	D' = 0.416

Haplotype	Frequency	Expectation under LE

---------	---------	--------------------

GG	0.050	0.025

CG	0.241	0.266

GC	0.035	0.060

CC	0.674	0.648

In phase alleles are GG/CC
