## Supplementary Figures for "Machine Learning-GWAS reveals the role of *WSD1* gene for cuticular wax ester biosynthesis and key genomic regions controlling early maturity in bread wheat" (from another version of this preprint: version 1, 2023-11-05)

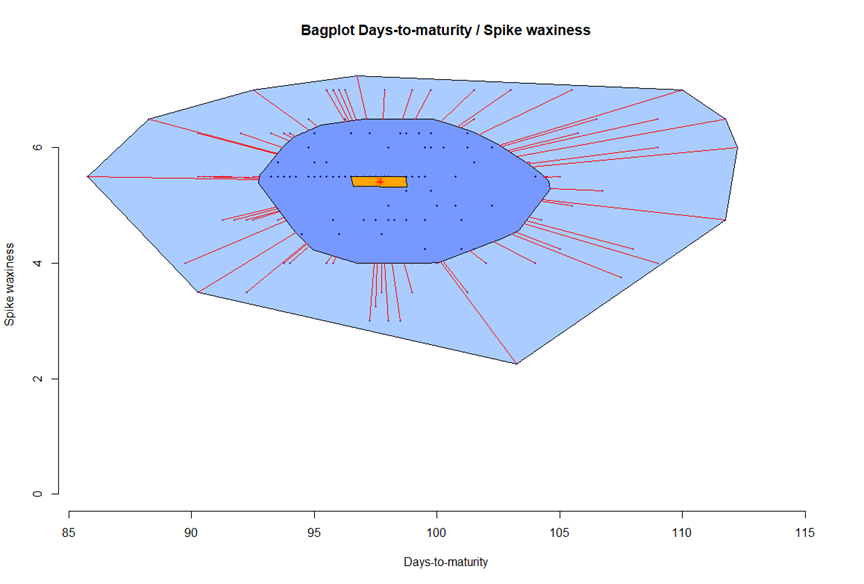

Supplementary Figure S1. Bagplot showing the relationship between spike waxiness and Days-to-Maturity traits in wheat varieties.

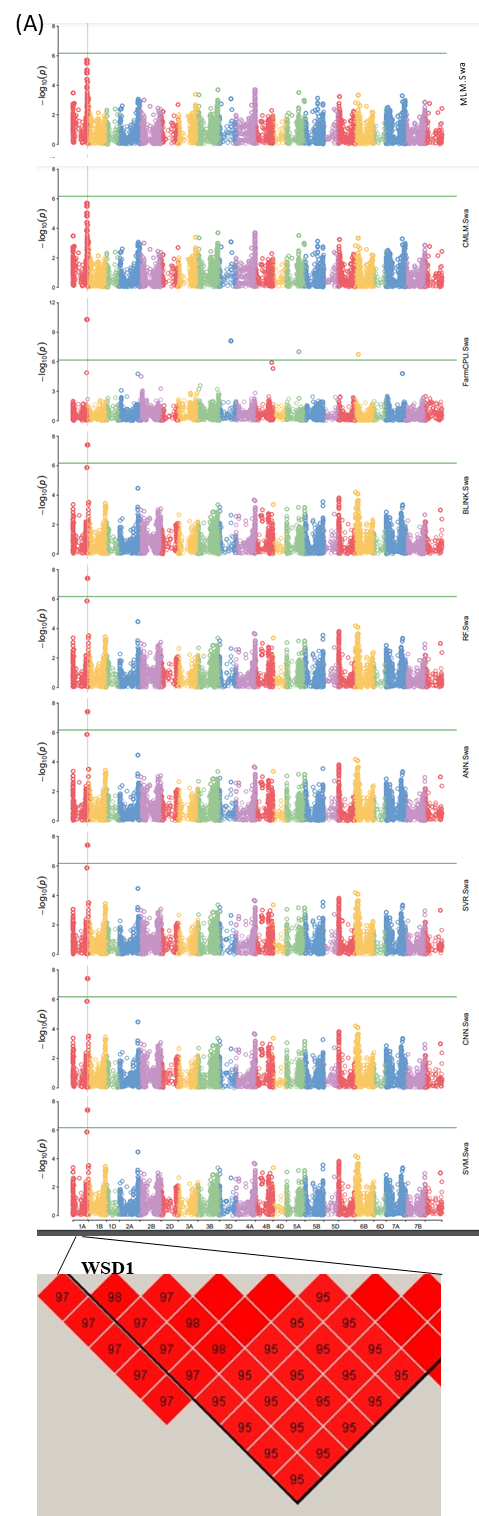

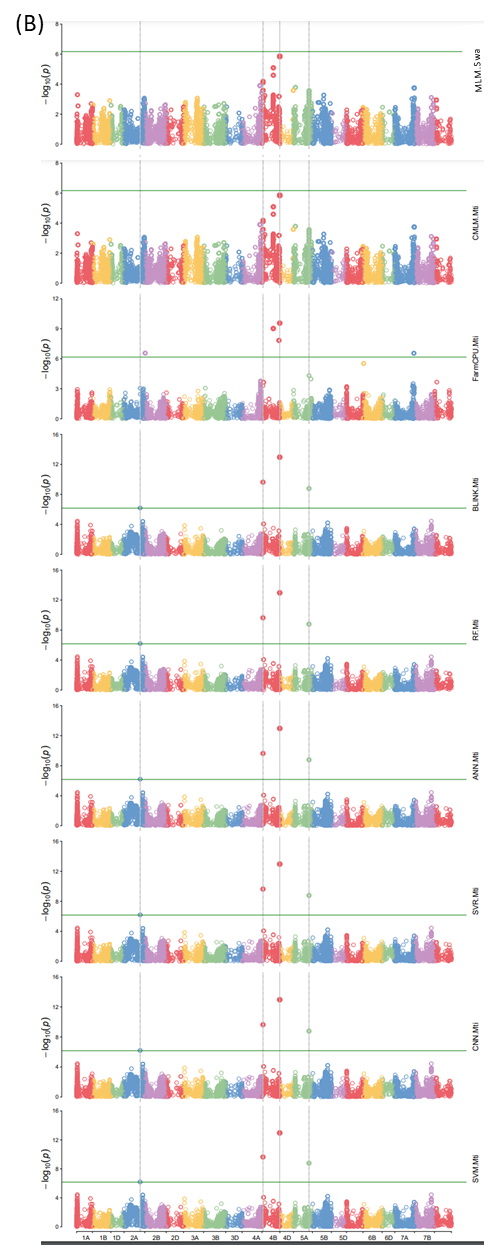

Supplementary Figure S2. Genomics regions involved in the regulation of SW and DM within the haplotype block containing the peak SNP for QTL on chromosome 1A (A) and chromosome 4B (B). respectively. Linkage disequilibrium plots showing regional Manhattan plot representing marker-trait associations associated with SW on chromosome 1A.
